# Estimating Ectopic Beat Probability with Simplified Statistical Models that Account for Experimental Uncertainty

**DOI:** 10.1101/2021.03.08.434403

**Authors:** Qingchu Jin, Joseph L. Greenstein, Raimond L. Winslow

## Abstract

Ectopic beats (EBs) are cellular arrhythmias that can trigger lethal arrhythmias. Simulations using biophysically-detailed cardiac myocyte models can reveal how model parameters influence the probability of these cellular arrhythmias, however such analyses can pose a huge computational burden. Here, we develop a simplified approach in which logistic regression models (LRMs) are used to define a mapping between the parameters of complex cell models and the probability of EBs (P(EB)). As an example, in this study, we build an LRM for P(EB) as a function of diastolic cytosolic Ca^2+^ concentration ([Ca^2+^]_i_), sarcoplasmic reticulum (SR) Ca^2+^ load, and kinetic parameters of the inward rectifier K^+^ current (I_K1_) and ryanodine receptor (RyR). This approach, which we refer to as arrhythmia sensitivity analysis, allows for evaluation of the relationship between these arrhythmic event probabilities and their associated parameters. This LRM is also used to demonstrate how uncertainties in experimentally measured values determine the uncertainty in P(EB). In a study of the role of [Ca^2+^]_SR_ uncertainty, we show a special property of the uncertainty in P(EB), where with increasing [Ca^2+^]_SR_ uncertainty, P(EB) uncertainty first increases and then decreases. Lastly, we demonstrate that I_K1_ suppression, at the level that occurs in heart failure myocytes, increases P(EB).

**Author summary:** An ectopic beat is an abnormal cellular electrical event which can trigger dangerous arrhythmias in the heart. Complex biophysical models of the cardiac myocyte can be used to reveal how cell properties affect the probability of ectopic beats. However, such analyses can pose a huge computational burden. We develop a simplified approach that enables a highly complex biophysical model to be reduced to a rather simple statistical model from which the functional relationship between myocyte model parameters and the probability of an ectopic beat is determined. We refer to this approach as arrhythmia sensitivity analysis. Given the efficiency of our approach, we also use it to demonstrate how uncertainties in experimentally measured myocyte model parameters determine the uncertainty in ectopic beat probability. We find that, with increasing model parameter uncertainty, the uncertainty in probability of ectopic beat first increases and then decreases. In general, our approach can efficiently analyze the relationship between cardiac myocyte parameters and the probability of ectopic beats and can be used to study how uncertainty of these cardiac myocyte parameters influences the ectopic beat probability.

## Introduction

Delayed after-depolarizations (DADs) are spontaneous depolarizations that occur during diastole [1]. When their amplitude is sufficiently large, DADs can trigger action potentials (APs) known as ectopic beats (EBs) [2]. Under certain conditions, for example in the setting of catecholaminergic polymorphic ventricular tachycardia, DADs can also trigger large scale cardiac arrhythmias[3]. Computational modeling of EBs and DADs has provided insights into the mechanisms by which they can trigger arrhythmias in the heart [4, 5].

We seek to understand how variations in the underlying biophysical properties or physiological state of the myocyte influence the probability of occurrence of EBs. To do this, we use a three-dimensional spatial model of the ventricular myocyte developed by Walker et al [5] in which the fundamental events governing intracellular calcium (Ca^2+^) dynamics are modeled stochastically. This model has previously been shown to reproduce realistic Ca^2+^ waves, DADs, and ectopic beats driven by stochastic gating of L-type Ca^2+^ channels (LCCs) and sarcoplasmic reticulum (SR) Ca^2+^ release channels (ryanodine receptors, RyRs).

The complexity of this stochastic model makes it challenging to perform the many repeated simulations needed to estimate the probability of ectopic beats, denoted P(EB), as a function of underlying model parameters. Here we explore an approach designed for estimating P(EB) that is computationally efficient. To do this we leverage a technique first developed by Sobie et al [6, 7] and apply it to formulate a logistic regression model (LRM) that maps myocyte model inputs (MMIs), which refers collectively to myocyte model parameters as well as state variable initial conditions, to probabilities of cellular arrhythmias. This approach, which we refer to as arrhythmia sensitivity analysis, allows for evaluation of the relationship between these event probabilities and their associated MMIs.

Use of an LRM to map MMIs to the probability of cellular arrhythmias also enables analyses of how experimental uncertainty in the measurement of MMIs influences estimates of these probabilities. One mainstream method for experimental uncertainty analysis is Monte Carlo simulation [8]. It has been used extensively in system biology and myocyte modeling [9-11]. However, such methods require many repeated model simulations, an approach which can become computationally demanding when using complex models of the myocyte such as that of Walker et al. [5]. LRMs also provide a computationally efficient approach for performing uncertainty analyses.

In this study, we develop a LRM model, mapping 4 MMIs (diastolic cytosolic Ca^2+^ concentration ([Ca^2+^]_i_), SR Ca^2+^ load ([Ca^2+^]_SR_), conductance of the inward rectifier current I_K1_ (G_K1_), and RyR opening rate (k_RyR_^+^)) to P(EB). We also investigate the role of uncertainty in [Ca^2+^]_SR_ on P(EB). It surprisingly reveals that as [Ca^2+^]_SR_ uncertainty increases, P(EB) uncertainty first increases (from a sharp unimodal distribution to a broad uniform-like distribution) and then decreases (toward a bimodal distribution with P(EB) concentrated at 0 or 1). Investigation of G_K1_ uncertainty showed that P(EB), with G_K1_ at reduced levels associated with heart failure, is significantly higher than that at normal G_K1_ density.

## Results

### Modeling Ectopic Beat Probability

An LRM was formulated to capture the quantitative relationship between P(EB) and 4 MMIs ([Ca^2+^]_i_, [Ca^2+^]_SR_, G_K1_ scaling factor (G_K1__sf) and k_RyR_^+^ scaling factor (k_RyR_^+^_sf)), where [Ca^2+^]_i_ and [Ca^2+^]_SR_ are the initial values of cytosolic and SR Ca^2+^ concentration, respectively, for each model simulation (i.e. realization). Because of the stochastic nature of the myocyte model, each realization may or may not generate a DAD or an ectopic beat. We defined a biophysically meaningful region of interest over which each of these parameters are varied and within which the LRM is built (S1 Table). Fig 1A shows 5 examples each of myocyte model simulated ectopic beats and DADs. Since ectopic beats consistently exhibit membrane potential waveforms that exceed 0 mV whereas DADs do not, a threshold of 0 mV was used to detect such events in each realization. Fig 1B demonstrates the relationship between this region of interest and P(EB) for both the underlying myocyte model and the LRM, where <u>B</u> ^T^ <u>P</u> (the argument of the logistic regression function) is the weighted (<u>B</u>) summation of LRM features (<u>P</u>).

**Fig 1.**
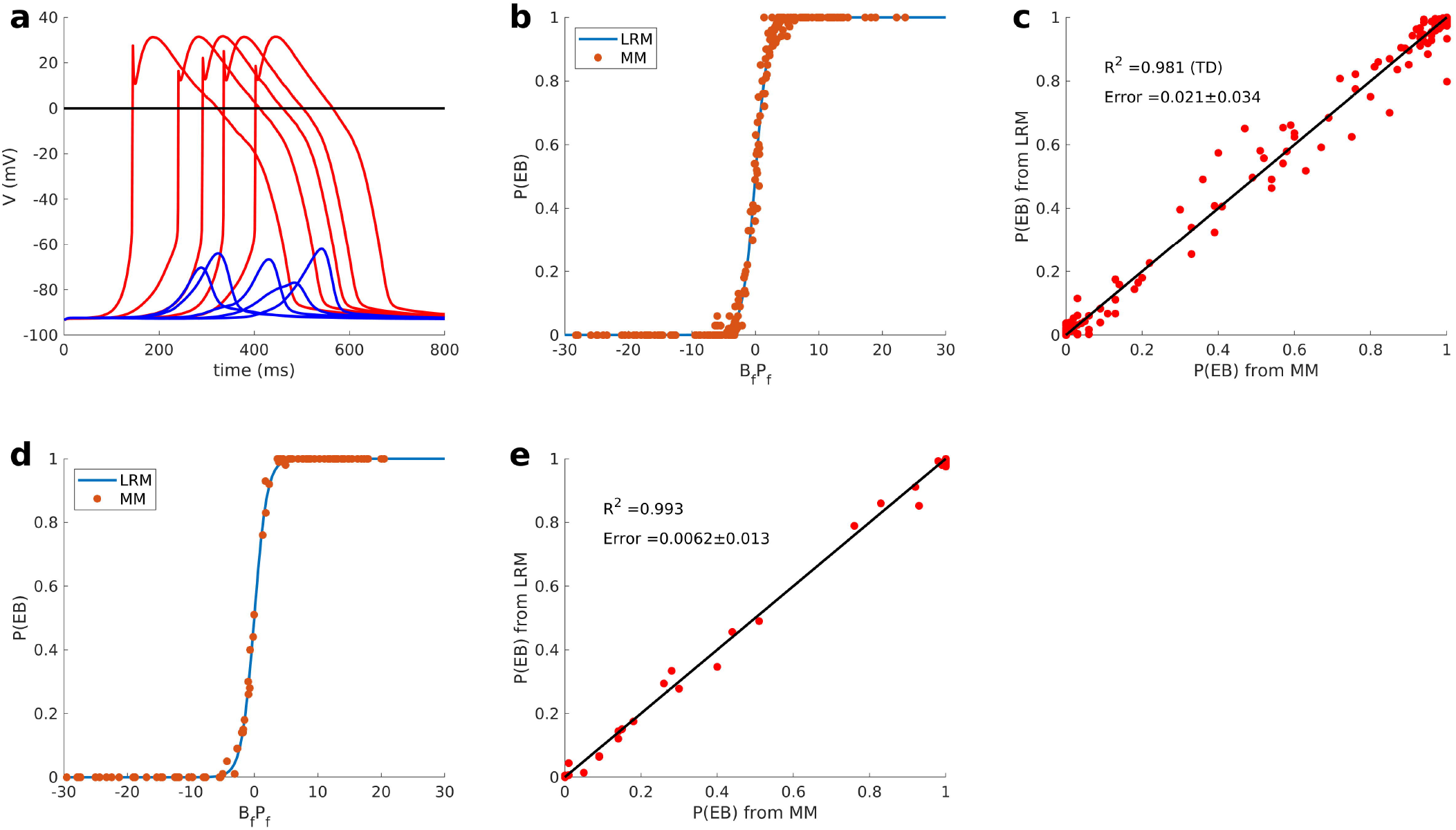
Ectopic beat study. (A) Simulations of ectopic beats (red) and simulations that generate DADs without induced ectopic beats (blue). (B) Comparison between LRM- predicted P(EB)s (blue) and myocyte model-generated actual P(EB)s (red) as a function of <u>B</u>^T^<u>P</u>, the argument of the logistic equation (S3 Eq for the training set). (C) Direct comparison between LRM-predicted P(EB)s (y-axis) and myocyte model-generated actual P(EB)s (x-axis) for the training set. (D) and (E) show the same analyses described in (B) and (C), respectively, for the independent test set. R^2^ is calculated only from MMI sets within the transition domain, LRM: logistic regression model, MM: myocyte model

The nonlinear logistic relationship defines three domains of interest: transition domain, lower plateau domain, and upper plateau domain. The transition domain is a region in which P(EB) is a sensitive function of <u>B</u>^T^<u>P</u>, corresponding to the steep part of the curve in Fig 1B. The lower plateau domain and upper plateau domain are regions where P(EB) is relatively insensitive to MMI perturbation, with P(EB) = ∼0 in the lower plateau domain and P(EB) = ∼1 in the upper plateau domain. To build the LRM, we sampled 200 MMI sets from the region of interest and ran 100 realizations for each set from which we obtain P(EB). Since the transition domain is relatively narrow and P(EB) is a very steep function within it, we have developed a two-stage sampling strategy to ensure adequate sampling of this domain (see Methods and S1 Text for details). The 200 MMI sets and their corresponding P(EB)s were used to train the LRM. Once determined, the LRM enables direct calculation of P(EB) for any MMI set without the need to simulate the underlying complex stochastic three-dimensional myocyte model. Fig 1B shows the fidelity with which the LRM model (blue line) reproduces P(EAD) estimated using the myocyte model (red points) for the training data set (200 MMI sets) as a function of <u>B</u>^T^<u>P</u>. Fig 1C shows the predicted P(EB) from the LRM versus P(EB) computed from the myocyte model simulations. The LRM performs well in predicting P(EB) (R^2^ = 0.981 for the 102 MMI sets inside the transition domain, mean prediction error of 0.021 ± 0.034 (see Methods and Eq 2)), despite the highly complex nonlinear properties of the myocyte model. To test the ability of the LRM to generalize, Figs 1D- E compare the LRM predicted P(EB)s to P(EB)s computed using the myocyte model for test data consisting of 100 independent MMI sets. The LRM performs well on the test set (R^2^ = 0.993 for the 22 sets within the transition domain, mean prediction error = 0.006 ± 0.013). Detailed modeling methods and performance metrics are described in Methods and S1 Text.

The LRM for P(EB) was formulated with a total of 10 features. Four of these are linear features: [Ca^2+^]_i_, [Ca^2+^]_SR_, Gk1_sf, and k_RyR_^+^_sf. In addition, the following 6 derived quadratic features were used: [Ca^2+^]_SR_^2^, [Ca^2+^]SR*k_RyR_^+^_sf, k_RyR_^+^_sf^2^, G_K1__sf^2^, [Ca^2+^]_i_^2^, and G_K1__sf*k_RyR_ ^+^_sf. Prior to fitting the LRM, features were scaled to a range of 0 to 1 in order to ensure interpretability of the LRM weights. Table 1 shows the LRM weights ranked by feature importance. Feature [Ca^2+^]SR is most important and is followed by feature k_RyR_ ^+^. [Ca^2+^]_SR_ ^2^ ranks as the third most important feature, indicating that quadratic features play an important role in improving model performance. In S2B Fig, the performance of the LRM built without quadratic features on the same test set has lower R^2^ (0.862) in the transition domain and higher error (0.023 ± 0.058) than the 10-feature model shown in S2D Fig. This confirms that the additional 6 quadratic features improve LRM performance.

**Table 1.** Ectopic beat features ranked by relative importance.

| Ranking | Features | Weights ± SE<br>from LRM |
| --- | --- | --- |
| 1 | $[Ca^{2+}]_{SR}$ | $71.27 \pm 3.81$ |
| 2 | $k_{RyR}^+_{sf}$ | $44.37 \pm 2.56$ |
| 3 | $[Ca^{2+}]_{SR}^2$ | $-24.41 \pm 2.13$ |
| 4 | $[Ca^{2+}]_{SR} * k_{RyR}^+_{sf}$ | $-15.36 \pm 2.76$ |
| 5 | $k_{RyR}^+_{sf}^2$ | $-13.45 \pm 1.14$ |
| 6 | $[Ca^{2+}]_i$ | $12.77 \pm 0.53$ |
| 7 | $G_{K1\_sf}$ | $-7.98 \pm 0.62$ |
| 8 | $G_{K1\_sf}^2$ | $-3.62 \pm 0.49$ |
| 9 | $[Ca^{2+}]_i^2$ | $-2.68 \pm 0.46$ |
| 10 | $G_{K1\_sf} * k_{RyR}^+_{sf}$ | $-2.48 \pm 0.54$ |

These results presented here demonstrate that LRMs can accurately predict P(EB) generated by the complex myocyte model. For the 100 MMI test sets used in each case, thousands of CPU hours are required to obtain event probabilities from the myocyte model whereas evaluation of LRMs requires negligible computing time on the order of milliseconds. The computationally demanding task of estimating these probabilities is made possible by this LRM approach.

In Figs 2A-D, P(EB) increases with increasing [Ca^2+^]_i_ or [Ca^2+^]_SR_. I_K1_ downregulation also increases P(EB). These findings agree with earlier simulations by Walker et al. [5]. The positive correlation between k_RyR_ ^+^_sf and P(EB) agrees with the finding of Paavola et al. that k_RyR_ ^+^ is positively related to DAD induction [12]. Figs 2E and F show multiple myocyte model realizations for two example MMI sets. Fig 2E is an example with relatively low P(EB) obtained using the set: [Ca^2+^]_i_ = 216 nM, [Ca^2+^]_SR_ = 0.47 mM, G_K1__sf = 0.699, and k_RyR_ ^+^_sf = 5.60, where 19 out of 100 simulations generated ectopic beats. Fig 2F is an example with relatively high P(EB) obtained using the set: [Ca^2+^]_i_=154 nM, [Ca^2+^]_SR_=0.60 mM, G_K1__sf = 0.775, and k_RyR_ ^+^_sf = 5.13, where 92 out 100 simulations generated ectopic beats. The only difference across simulations in either Fig 2E or 2F is the random gating of LCCs and RyRs.

**Fig 2.**
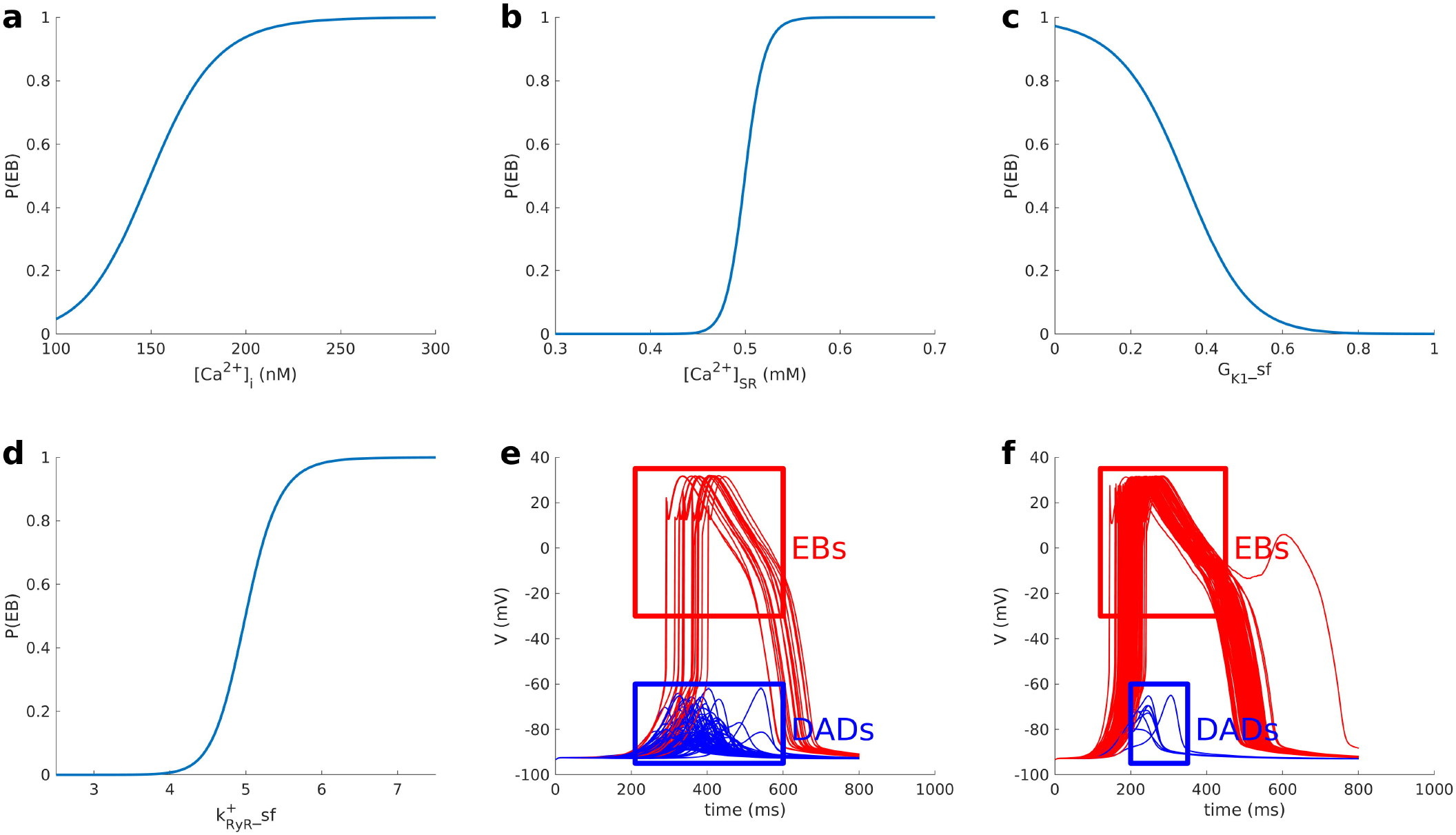
Relationship between MMIs and P(EB). In A-D, with exception of the MMI that is varied along the x-axis, MMIs are fixed at: [Ca^2+^]_i_ =150 nM, [Ca^2+^]_SR_ = 0.5 mM, G_K1__sf = 0.338, and k_RyR_^+^_sf = 5. (E) 100 random myocyte model simulations with a low P(EB) = 0.19; MMI set: [Ca^2+^]_i_= 216 nM, [Ca^2+^]_SR_ = 0.47 mM, G_K1__sf = 0.699, and k_RyR_ ^+^_sf = 5.60. (F) 100 random myocyte model simulations with a high P(EB) = 0.92; MMI set: [Ca^2+^]_i_ =154 nM, [Ca^2+^]_SR_ = 0.60 mM, G_K1__sf = 0.775, and k_RyR_^+^_sf = 5.13. Red curves represent EBs and blue curves represent DADs that do not trigger ectopic beats.

### Uncertainty analysis of P(EB)

The LRM allows for analysis of the uncertainty of probabilistic events such as ectopic beats arising from the experimental uncertainties inherent in measurements of the MMIs. We treat MMIs as random variables, and then the LRM becomes a probability transformation function (i.e. S3 Eq) of these MMI random variables. MMIs with uncertainties are assumed to have a normal distribution with σ taking the experimentally measured value for each MMI. From the experimentally derived distribution for each MMI, we randomly generate 10^6^ sets of MMIs and evaluate the LRM for each set to generate a predicted distribution of P(EB).

In the following study, the effect of [Ca^2+^]_SR_ uncertainty on P(EB) was determined. In Figs 3A-C, MMI set 1 was assumed to represent mean values for each MMI: [Ca^2+^]_i_ = 150nM, [Ca^2+^]_SR_ = 500µM, k _RyR_^+^_sf = 5, and G_K1__sf = 0.338, which yields P(EB) = 0.5. We assumed three different degrees of uncertainty in [Ca^2+^]_SR_ corresponding to σ values of 5 µM, 15 µM, and 30 µM. In Figs 3A-C, histograms show the P(EB) distribution given different [Ca^2+^]_SR_ uncertainties, which leads to 3 different distribution patterns: unimodal (Fig 3A), approximately uniform (Fig 3B), and bimodal (Fig 3C). The red line indicates the mean P(EB). Fig 3A (σ = 5 µM) is a unimodal distribution in which P(EB) takes on values at or near 0.5 with high probability. Fig 3B (σ = 15 µM) is an approximately uniform distribution between 0 and 1. Thus, at this level of uncertainty the distribution of P(EB) has maximum entropy. In Fig 3C (σ = 30 µM), the distribution of P(EB) becomes bimodal, with two peaks located at P(EB) values close to 0 and 1. In this scenario, P(EB) is equally likely to be either 0 or 1 given the symmetry of the P(EB) distribution.

**Fig 3.**
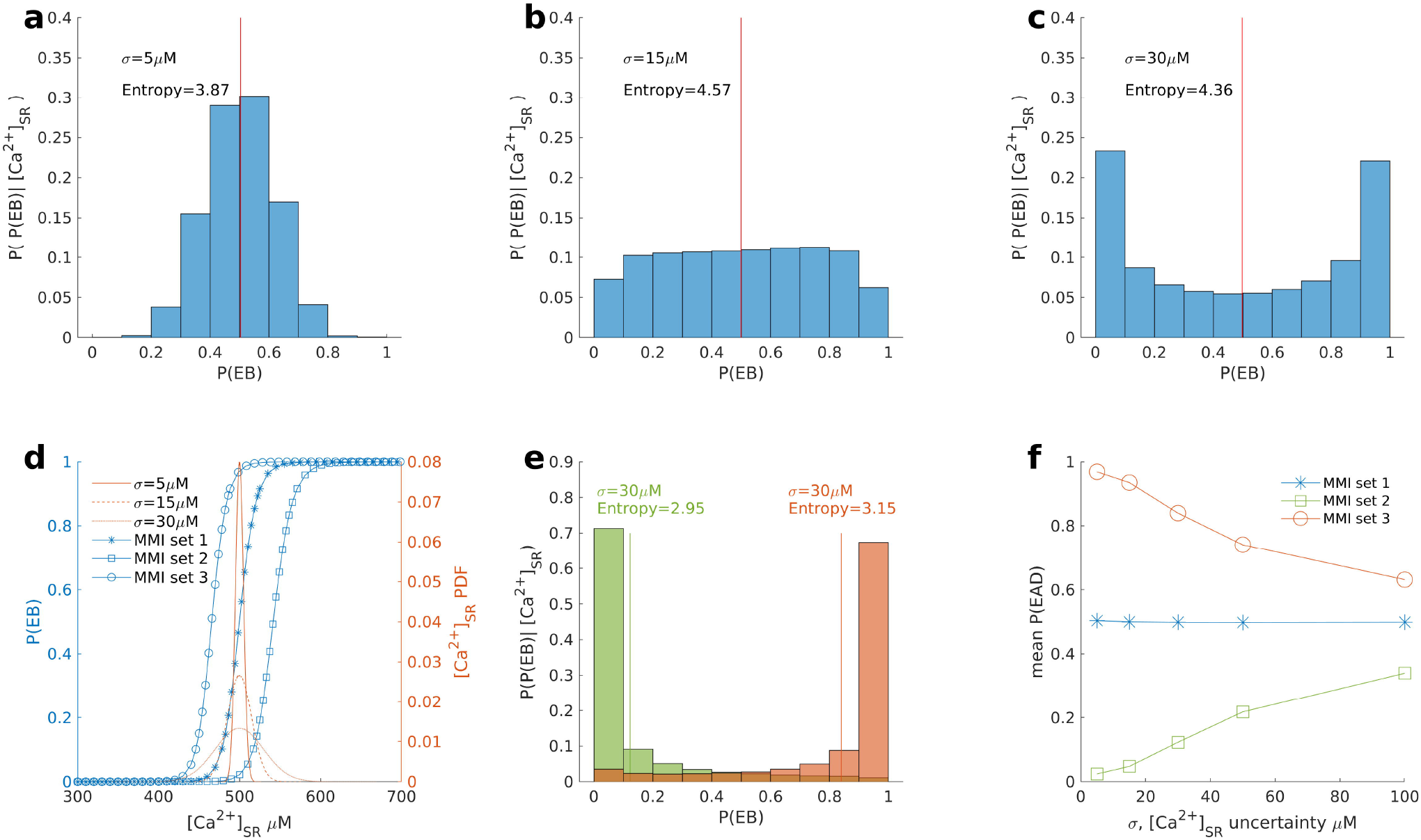
Uncertainty analysis of P(EB) due to variation in [Ca^2+^]_SR_. Three MMI sets are used. All MMI sets have [Ca^2+^]_i_ = 150 nM, [Ca^2+^]_SR_ = 500 µM, and k_RyR_ ^+^_sf = 5. G_K1__sf for set 1, set 2, and set 3 is 0.338, 0.638, and 0.000, respectively. The P(EB) distributions given different hypothetical [Ca^2+^]_SR_ uncertainties (σ) of set 1, where (A) σ = 5 µM, (B) σ = 15 µM, and (C) σ = 30 µM. The vertical red line is the mean of the P(EB) distribution. P(P(EB)|[Ca^2+^]_SR_), on the y-axis, refers to the P(EB) distribution given [Ca^2+^]_SR_ uncertainty. (D) Red curves are the [Ca^2+^]_SR_ PDFs for σ = 5µM (solid line), σ = 15 µM (dashed line), and σ = 30 µM (dotted line). Blue curves are the model characteristic (MC) curves relating [Ca^2+^]_SR_ and P(EB) for set 1 (stars), set 2 (open squares), and set 3 (open circles). (E) P(EB) distributions with [Ca^2+^]_SR_ uncertainty σ = 30 µM for set 2 (green bars) and set 3 (red bars). (F) Relationship between mean P(EB) and [Ca^2+^]_SR_ uncertainty for all three sets.

In Fig 3D, blue curves show the [Ca^2+^]_SR_-P(EB) relationship determined by the LRM, which we refer to as the model characteristic (MC) curve for [Ca^+^]_SR_, and the red curves show [Ca^2+^]_SR_ probability density functions ([Ca^2+^]_SR_ PDF) which widens as σ increases. A comparison of the relative positions of the MC curve and the [Ca^2+^]_SR_ PDF can be used to explain the previously described P(EB) distribution patterns. The MC curve for set 1 (stars) shows the [Ca^2+^]_SR_-P(EB) relationship as [Ca^2+^]_SR_ is varied. The width of the [Ca^2+^]_SR_ PDF for σ = 5 µM (solid curve) is quite narrow and exists at the center of the transition domain where P(EB) = 0.5. As a result, the distribution of P(EB) is unimodal with peak near 0.5. The width of the [Ca^2+^]_SR_ PDF for σ = 15 µM (dashed curve) coincides with the boundaries of the transition domain such that the distribution of P(EB) in Fig 3B is approximately uniform. The [Ca^2+^]_SR_ PDF for σ = 30µM (dotted line) is much wider such that a sufficiently large portion of the [Ca^2+^]_SR_ PDF coincides with the lower plateau domain and the upper plateau domain of the MMI set 1 MC curve, where P(EB) takes on values of either 0 or 1 resulting in a bimodal distribution.

This behavior reveals an interesting finding that with increasing uncertainty of [Ca^2+^]_SR_, uncertainty in P(EB) first increases and then decreases. To confirm our finding, we calculated the entropy of P(EB) distributions for Figs 3A-C by S4 Eq. The P(EB) distribution with σ = 15 µM has the highest entropy which validates our finding. The mean P(EB)s for all distributions shown in Figs 3A-C are 0.5 because all variations in [Ca^2+^]_SR_ occur about a mean [Ca^2+^]_SR_ = 500 µM in this scenario.

The role of [Ca^2+^]_SR_ uncertainty on P(EB) is evaluated for two additional sets for which P(EB) ≠ 0.5: set 2 ([Ca^2+^]_i_= 150 nM, [Ca^2+^]_SR_ = 500 µM, k_RyR_^+^_sf = 5, and G_K1__sf = 0.638, for which P(EB) = 0.02) and set 3 ([Ca^2+^]_i_ = 150 nM, [Ca^2+^]_SR_ = 500 µM, k_RyR_^+^_sf = 5, and G_K1__sf = 0.0, for which P(EB) = 0.97). The distribution of P(EB) for sets 2 and 3 are evaluated for [Ca^2+^]_SR_ uncertainty with σ = 30µM. In Fig 3E, the P(EB) distribution of [Ca^2+^]_SR_ is unimodal for both set 2 and set 3. MC curves for both set 2 (open squares) and 3 (open circles) are shown in Fig 3D. For set 2, the [Ca^2+^]_SR_ PDF (σ = 30 µM) coincides with the lower plateau domain and part of the transition domain of the MC curve. This results in a P(EB) distribution that is a unimodal with P(EB) values close to 0. For set 3, the MC curve is shifted such that the [Ca^2+^]_SR_ PDF (σ = 30 µM) coincides with the upper plateau domain and part of the transition domain of the MC curve. This results in a P(EB) distribution that is a unimodal with P(EB) values close to 1. These demonstrate that the P(EB) distribution pattern depends on the width of [Ca^2+^]_SR_ PDF (degree of [Ca^2+^]_SR_ uncertainty) as well as the relative position of the mean [Ca^2+^]_SR_ with respect to the transition domain of its MC curve.

In general, [Ca^2+^]_SR_ uncertainty impacts the mean of P(EB). Fig 3F shows the relationship of [Ca^2+^]_SR_ uncertainty (σ) and the mean P(EB) for sets 1-3. Experimental uncertainty in measurements of [Ca^2+^]_SR_ varies widely from 10 µM to 290 µM [13-15], and we therefore evaluated 0 µM ≤ σ ≤ 100 µM. Results show that with the increase of [Ca^2+^]_SR_ uncertainty, the mean P(EB) converges to 0.5 regardless of the selected set. When the [Ca^2+^]_SR_ PDF is narrow (e.g., σ = 5µM), the P(EB) distribution will be unimodal centered at the value of P(EB) defined by the LRM for the given set, and therefore, the mean P(EB) will be close to this value. In contrast, when the [Ca^2+^]_SR_ uncertainty is sufficiently large, a majority of the [Ca^2+^]_SR_ PDF will coincide with the lower plateau domain and upper plateau domain of the MC curve, with only a small to negligible portion of the [Ca^2+^]_SR_ PDF coinciding with the transition domain of the MC curve. In the limit as σ grows to be large, for any set, the distribution of [Ca^2+^]_SR_ flattens such that there is equal probability of it taking on values in either the lower plateau domain or upper plateau domain of the MC curve, and hence P(EB) becomes equally likely to take on values of 0 or 1 and mean P(EB) converges to 0.5 (Fig 3F).

### Role of G_K1_ on P(EB)

The effect of I_K1_ current density scaling factor (G_K1__sf) uncertainty on P(EB) was evaluated. We used the experimental results of Pogwizd et. al. [16] to obtain the distribution of G_K1__sf values in both normal and heart failure (HF) myocytes. The process by which the LRM parameters (i.e. weights) are constrained is described in detail in the supplement (S3A and S3B Figs). The following three MMIs are fixed as follows: [Ca^2+^]_i_ = 150 nM, [Ca^2+^]_SR_ = 550 µM, and k_RyR_^+^_sf = 5. G_K1__sf is treated as uncertain where in normal cells it is 1.0000 ± 0.1108 (σ_N_) and in heart failure (HF) cells it is 0.5100 ± 0.0852 (σ_HF_). The P(EB) corresponding to the mean value of G_K1__sf is 0.008 and 0.918 for normal and HF, respectively. Fig 4A shows the P(EB) distributions arising from the experimental uncertainty of G_K1__sf. The P(EB) arising from the experimental distribution of G_K1__sf is 0.0294 ± 0.0611 in normal cells and 0.8788 ± 0.1183 in HF. Results show that, even when accounting for experimental uncertainty, P(EB) for HF G_K1__sf is significantly higher than that corresponding to normal G_K1__sf. Fig. 4B shows the relative positions of the G_K1__sf MC curve (blue) and the G_K1__sf PDFs (red). The normal and HF G_K1__sf PDFs exhibit very little overlap. The normal G_K1__sf PDF primarily coincides with a region of the MC curve where P(EB) < ∼0.3. In contrast, the HF G_K1__sf PDF primarily coincides with a region of the MC curve where P(EB) > ∼0.4. The relationships between the G_K1__sf PDFs and the MC curve lead to the distributions of P(EB) in Fig 4A.

**Fig 4.**
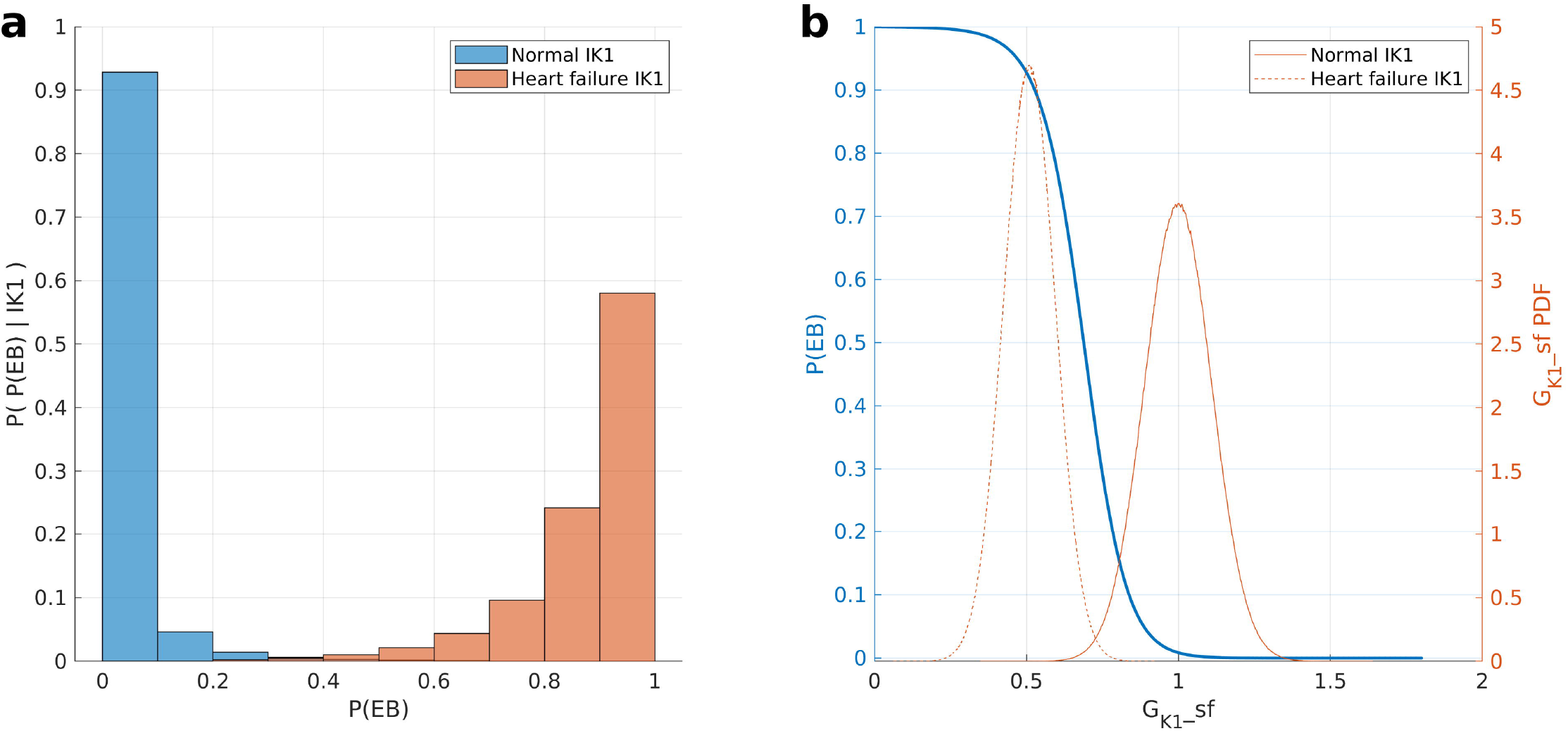
Effect of G_K1__sf uncertainty on P(EB). (A) Distributions of P(EB) for normal G_K1__sf (blue) and HF G_K1__sf (red). (B). MC curve (blue) shows the relationship of G_K1__sf and P(EB) for the LRM. G_K1__sf PDFs for normal (solid red line) and HF (dashed red line).

## Discussion

Regression models have been used previously by Sobie et al. to estimate action potential duration, peak and resting membrane potential, and Ca^2+^ transient amplitude from deterministic myocyte models, as well as the probability of the occurrence of Ca^2+^ sparks from a stochastic Ca^2+^ release site model [6, 7]. Based on their method, we developed a new pipeline for establishing simplified models (LRMs) for estimating the probability of cellular cardiac arrhythmia events simulated by a complex stochastic myocyte model. Fig 1E shows excellent prediction performance of the LRM in predicting the probability of ectopic beats. The method developed in this work is universal in the sense that it should be considered a general approach to simplifying highly complex nonlinear stochastic models in a wide range of applications.

Our approach differs from that of Lee at al. [7] at two points in the modeling pipeline. First, we developed a two-iteration strategy for generating MMI sets. The second iteration ensures that the transition domain is adequately sampled. Second, we derived LRM quadratic features from the MMIs. The rationale for this approach is that since the myocyte model is highly nonlinear, we expect improved performance by allowing for a nonlinear relationship between MMIs and P(EB) in the LRM. To demonstrate the performance improvement of these two steps, we built preliminary LRMs from only the first iteration MMI sets and using only MMIs as features (i.e. without derived quadratic features). For prediction of P(EB), this approach yields an average error of 0.026 ± 0.058 (S2A Fig), which is worse than that (0.006 ± 0.013) obtained with the full pipeline (S2D Fig).

The LRM approach has the benefit of computational efficiency. The average computational time required for each MMI set in the EB study is ∼0.4 (hours per realization) * 100 (realizations) = 40 CPU hours. To build the LRM, however, 200 training MMI sets must be simulated with the myocyte model, which takes ∼8000 CPU hours. After building the LRM, calculating for new MMI sets takes negligible time. For example, the computational time required to run the test set (100 MMI sets) in the ectopic beat study is therefore 4000 CPU hours. In contrast, the computational time for LRMs is negligible (∼5 CPU ms). Additionally, in the uncertainty study of [Ca^2+^]_SR_, 10^6^ samples were randomly sampled from the uncertainty range of [Ca^2+^]_SR_. If we estimate the P(EB) for each sample, the total computational time required is 0.4 * 100 * 10^6^ = 4*10^7^ CPU hour. In contrast, LRM takes less than 1 CPU second. These results show the dramatic improvement in computational efficiency of the LRM.

LRMs enable us to quantitatively explore the relationship between MMIs and event probabilities. In the ectopic beat study, Fig. 2A and 2B show that P(EB) increases with increasing [Ca^2+^]_i_ and [Ca^2+^]_SR_, indicating that elevation of cellular Ca^2+^ (i.e. Ca^2+^ overload) tends to increase the likelihood of ectopic beats. This result is consistent with previous experimental and computational studies [5, 17]. Fig. 2C shows that increasing G_K1__sf reduces P(EB). I_K1_ downregulation has been shown to be associated with ventricular arrhythmias such as Anderson’s syndrome [18] and LQTS [19]. Increased k_RyR_ ^+^_sf is also shown in Fig. 2D to elevate P(EB). Kubalova et al. showed that heart failure myocytes exhibit elevated RyR open probability [20]. Paavola et. al. also demonstrated the positive correlation between RyR opening rate with DAD induction and Ca^2+^ waves [12].

In Fig. 3A-C, in which [Ca^2+^]_SR_ variation was explored, three P(EB) distribution patterns (unimodal, approximately uniform, and bimodal) were revealed when exploring the role of [Ca^2+^]_SR_ uncertainty on predictions of P(EB). This implies that with the increase of [Ca^2+^]_SR_, the uncertainty of P(EB) first increases and then decreases. This result is also confirmed by comparing the entropies of P(EB) distributions associated with different degrees of uncertainty in [Ca^2+^]_SR_. Reported experimental uncertainties (standard errors) of [Ca^2+^]_SR_ vary widely from 10µM to 290µM [13-15]. A consequence of this is that for the majority of the realistic uncertainty range (30 - 290 µM), myocytes are unlikely to be in the transition domain where both DADs and EBs occur with nonzero probability. Namely, with a small perturbation in [Ca^2+^]_SR_ relative to the uncertainty range, the most likely behavior of the cell would jump between always exhibiting DADs and always exhibiting EBs. This is consistent with the fact that the EB is a bifurcation phenomenon in the stochastic myocyte model. Other studies also showed that EBs arise as a result of a bifurcation [21].

In Fig 4A, the heart failure (HF) G_K1_ study shows that, given the experimental uncertainty, P(EB) associated with HF G_K1_ is statistically significantly higher than P(EB) of normal G_K1_. Consistent with our results, Maruyama et al. showed that suppressing I_K1_ considerably enhanced the DAD amplitude and P(EB) [24]. This result shows the potential in using the LRM-predicted probability of an arrhythmic event to test the propensity for arrhythmias resulting from variations in other ion currents. We chose to explore G_K1_ in HF myocytes because I_K1_ data were available, and this method can be used to explore the impact of variation in other MMIs as well. These results also demonstrate the potential role for P(EB) as a useful biomarker for arrhythmia predictions, particularly in mutations and diseases linked to DADs such as catecholaminergic polymorphic ventricular tachycardia [25]. Additionally, early afterdepolarization (EAD), another important type of cellular arrhythmia, can be predicted via LRM. Such an LRM may be used as a biomarker to predict the arrhythmogenic risk of EAD-related diseases such as long QT syndrome [26].

In summary, we developed a pipeline for building logistic regression models (LRMs) that predict the probability of cellular arrhythmias. These simplified models faithfully reproduce the relationship between parameters/inputs and probabilistic events as learned from the mechanistic biophysically-detailed stochastic myocyte models, but with far less computational burden. As far as we know, this is the first simplified model enabling quantitative investigation of determinants of the probability of cellular arrhythmias. This approach, which we refer to as arrhythmia sensitivity analysis, allows for systematic study of the relationship between these event probabilities and their associated myocyte model inputs. As an example, we built a LRM to study the relationship between P(EB) and 4 MMIs: [Ca^2+^]_i_, [Ca^2+^]_SR_, G_K1_ and kRyR_+_. The negligible computational burden of LRM allows us to perform uncertainty analysis. In the [Ca^2+^]_SR_ uncertainty study, we showed and rationalized an emergent property of the P(EB) distribution, where with increasing [Ca^2+^]_SR_ uncertainty, P(EB) uncertainty first increases and then decreases. Lastly, the heart failure G_K1_ study shows the potential value of our approach in revealing the determinants of arrhythmia probability, indicating the potential for this method to be used as a tool for arrhythmia risk prediction.

## Methods

### Modified Walker Ventricular Myocyte Model

This study uses a modified version of the Walker et al. stochastic ventricular myocyte model [5]. The Walker et al. model is a biophysical-detailed three-dimensional spatial stochastic model of a cardiac ventricular myocyte containing 25,000 Ca^2+^ release units (CRUs) (each representing a dyad) along with their sub-membrane compartments. A subset of CRUs (5400) was used for all simulations to accelerate the computation and all CRU Ca^2+^ fluxes were scaled to the total number of CRUs. This model has been shown to reproduce realistic Ca^2+^ waves as well as DADs. Chu et al. [27] recently developed a model of the Na^+^-Ca^2+^ exchanger (NCX) that includes allosteric regulation of NCX function by Ca^2+^. This NCX model was incorporated into the Walker et al. model. Following Chu et al. [27], NCX was localized 15% in dyad, 30% in the submembrane compartment, and 55% in the cytosol. In addition to these changes, NCX conductance was increased 20%, the time constant of Ca^2+^ flux from submembrane to cytosol was reduced by 20%, IKr conductance increased by 20%, SERCA pump rate decreased by 25%, and number of Ito2 channels reduced from 8 to 5 in each submembrane compartment to assure a more realistic canine APD (∼290 ms) and Ca^2+^ transient peak (∼800 nM), similar to that of the normal canine ventricular myocyte [28]. The model code is available on the Github (https://github.com/JHU-Winslow-Lab/Ectopic-beat-paper-code.git).

### Simulation Protocol for Ectopic Beat

In the ectopic beat studies, the initial state of the myocyte model corresponds to the diastolic state, where membrane potential is -93.4 mV and NCX is partially allosterically activated (∼20%-40%). Model simulations are performed under a β- adrenergic stimulation protocol, which is the same as previously described by Walker et al. [29], with the exception that RyR opening rate is increased 4-fold compared to its control value based on experimental data for heart failure [30] and to increase the propensity for EBs. No external current was applied, and simulation duration was 800ms.

## Statistical Modeling

Fig 5 illustrates the general workflow for building a logistic regression model (LRM) from an underlying biophysically-detailed myocyte model. A more detailed workflow (S1 Fig) and its mathematical development is provided in the supplement. The steps of the workflow in Fig 5 are performed twice to build the LRM. In the first iteration, relevant myocyte model inputs (MMIs), which refers collectively to myocyte model parameters as well as state variable initial conditions, and their ranges of variation (regions of interest) are identified. A region of interest is established to keep MMIs within biophysically relevant ranges. A detailed description of region of interest is in the next section. Multiple randomly selected MMI sets are generated by uniform sampling within this region of interest. For each sampled MMI set, multiple simulations (realizations) are performed using the modified Walker model. A binary indicator random variable is defined using the output of each simulation. This random variable indicates whether the cellular arrhythmia event of interest did or did not occur. For each MMI set, an event (occurrence of an ectopic beat) probability is estimated based on its frequency of occurrence over all realizations. All MMI sets and their corresponding event probabilities are then used to derive a logistic regression model relating the probability of the event to a weighted sum of MMIs:

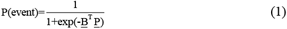

, where <u>P</u> = <u>P_o_</u> is a vector of features, <u>B</u> = <u>B_o_</u> is a vector of weights. The subscript “*o”* stands for “initial”, representing the first iteration of the pipeline. P(event) represents the estimated event occurrence probability.

**Fig 5.**
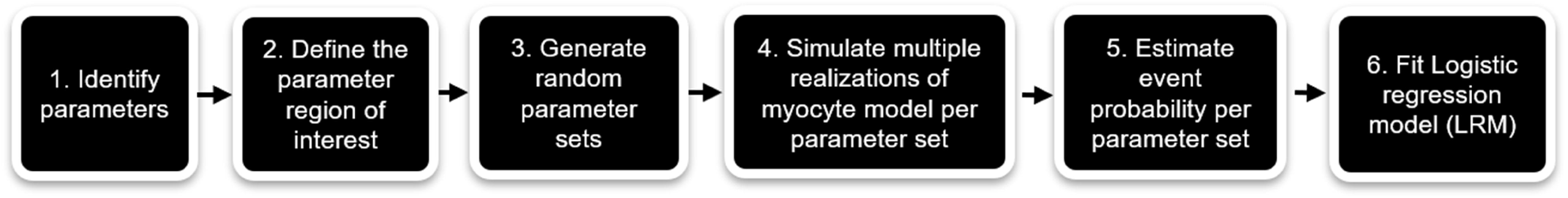
General workflow pipeline for logistic model formulation.

After this first iteration, the logistic equation (1) is then used as an additional constraint to estimate the transition domain, which is a subregion within the region of interest for the MMI set. A detailed description of transition domain is in the next section. The second iteration then samples only within the transition domain, and steps 4 – 5 of the workflow are repeated. MMI sets and corresponding event probabilities obtained from both iterations are combined, and the final LRM is built again using Eq 1, where <u>P</u> = <u>Pf</u> is the final feature vector and <u>B</u> = <u>Bf</u> is the final weights vector. The subscript “*f”* stands for “final”, representing the second iteration of the pipeline. We refer to this model as the LRM. 100 training MMI sets are generated in both the first and second iterations. For each MMI set, 100 realizations (for ectopic beat study) are simulated to estimate the probability of the event of interest. To ensure the generalizability of the LRM, an additional independent 100 test MMI sets are generated within the region of interest to be used for evaluation of the performance of the LRM. All logistic regression model fits were performed with MATLAB R2018b *fitglm* function.

### MMI Identification and MMI Region of Interest

When building the LRM for predicting probability of occurrence of DAD-induced EBs (P(EB)), four MMIs were selected to be varied. These were initial states of [Ca^2+^]_i_ and [Ca^2+^]_SR_, and scaling factors controlling I_K1_ current density (G_K1__sf) and RyR opening rate (kRyR^+^_sf). For simplicity, [Ca^2+^]_i_ and [Ca^2+^]_SR_ values mentioned in this paper are all initial state values. Upper and lower bounds on diastolic [Ca^2+^]_i_ and [Ca^2+^]_SR_ were obtained from experimental data [14, 31, 32]. Down-regulation of I_K1_ [16] and upregulation of RyR opening rate [20] have both been shown to occur in failing heart myocytes and changes in these cellular properties are associated with cellular arrhythmias.

The region of interest can be separated into two mutually exclusive subspaces in terms of the arrhythmia event probability: plateau domain and transition domain. The plateau domain is the space where MMI sets simulated in the myocyte model yield P(event) strictly equal to 0 or 1. The transition domain is the space where MMI sets simulated in the myocyte model yield P(event) strictly > 0 and < 1. We can specify the lower plateau domain as corresponding to P(event) = 0 and the upper plateau domain as corresponding to P(event) = 1 since these are also mutually exclusive. Because of the finite number of realizations performed with the myocyte model for each MMI set, P(event) takes on a discrete set of values such that we can strictly follow this definition.

### LRM Validation Metrics

Linear regression was performed between values of myocyte model-generated actual P(event) and LRM predicted P(event). The R-squared (R^2^) from the linear regression was calculated. Absolute error was also calculated as

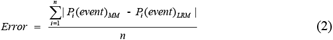

,where *n* is the number of MMI sets, P_i_(event)_MM_ is event probability of *i*th MMI set obtained from the myocyte model, and P_i_(event)_LRM_ is event probability of *i*th MMI set obtained from the LRM.

### Uncertainty Analyses

The LRM derived in this study is a function that maps MMIs to P(event). Experimental measurements of these MMIs exhibit variability, and this variation can be modeled by assuming these measurements are drawn from an underlying probability distribution. In this case, the event probability produced by the LRM is itself a random variable. We wish to characterize this uncertainty by computing the distribution of the estimated event probability. To do this, when the mean and variance of MMI estimates are available from experimental data, we assume MMIs are random variables drawn from a normal distribution with that mean and variance. In cases where the argument of the (<u>B</u>_f_ ^T^ <u>P</u>_f_) i involves a weighted sum of individual MMIs not including quadratic features, this distribution can be calculated analytically, and is the logit-normal distribution (Eq. S5) [33]. In cases where the feature vector <u>P</u>_f_ includes elements containing quadratic features, the distribution of P(event) must be estimated numerically. In this latter case, we randomly generate 10^6^ sample vectors for a specific MMI set based on the underlying MMI uncertainty distributions and use the LRM to predict P(event) for all sample vectors. Following this approach, the P(event) distribution can be estimated.

## Supporting information

S1 Text: Supporting description of logistic regression model and entropy

## Supporting information

**S1 Fig.**
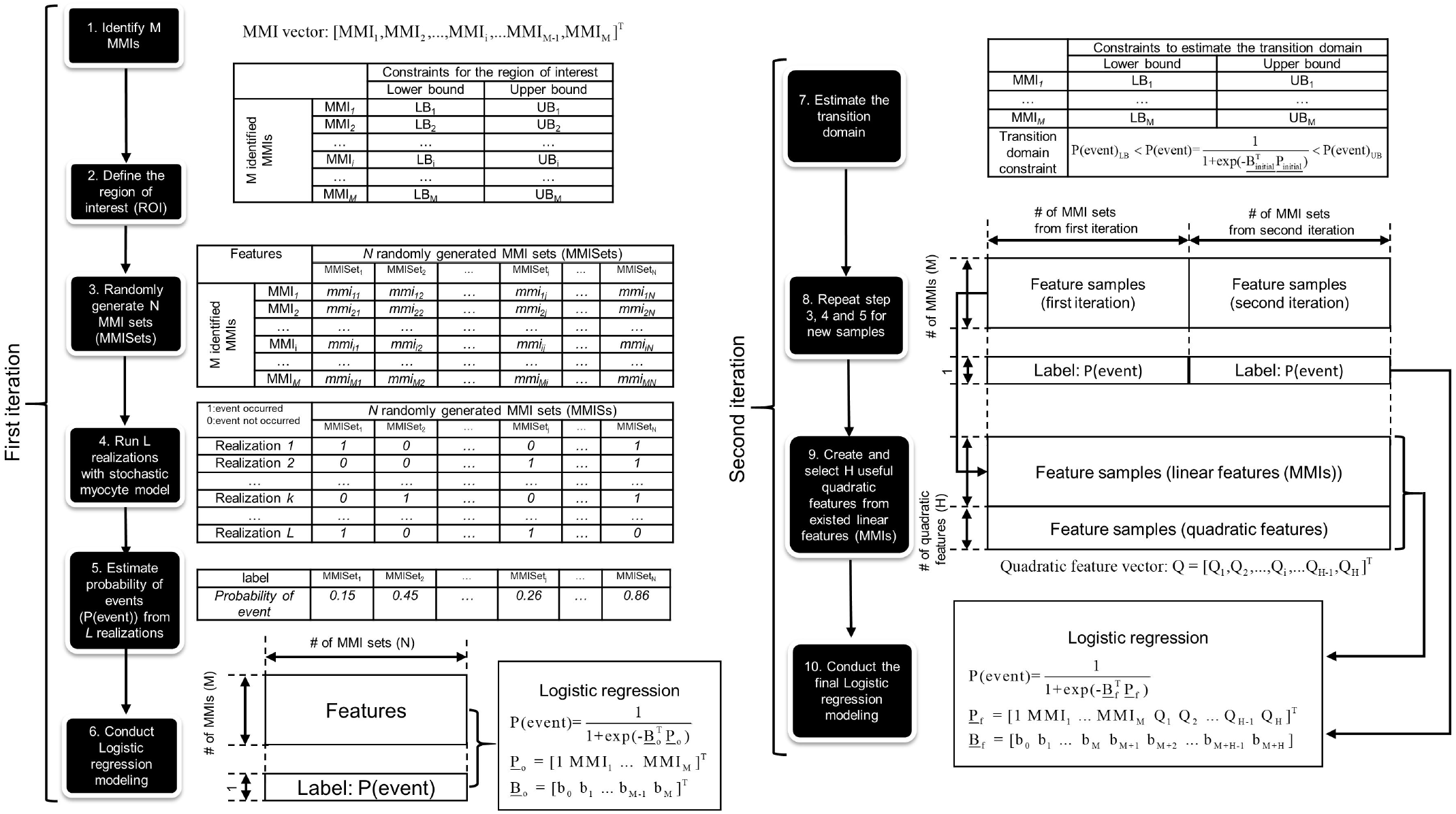
Detailed workflow for building the 2-iteration logistic regression model (LRM).

**S2 Fig.**
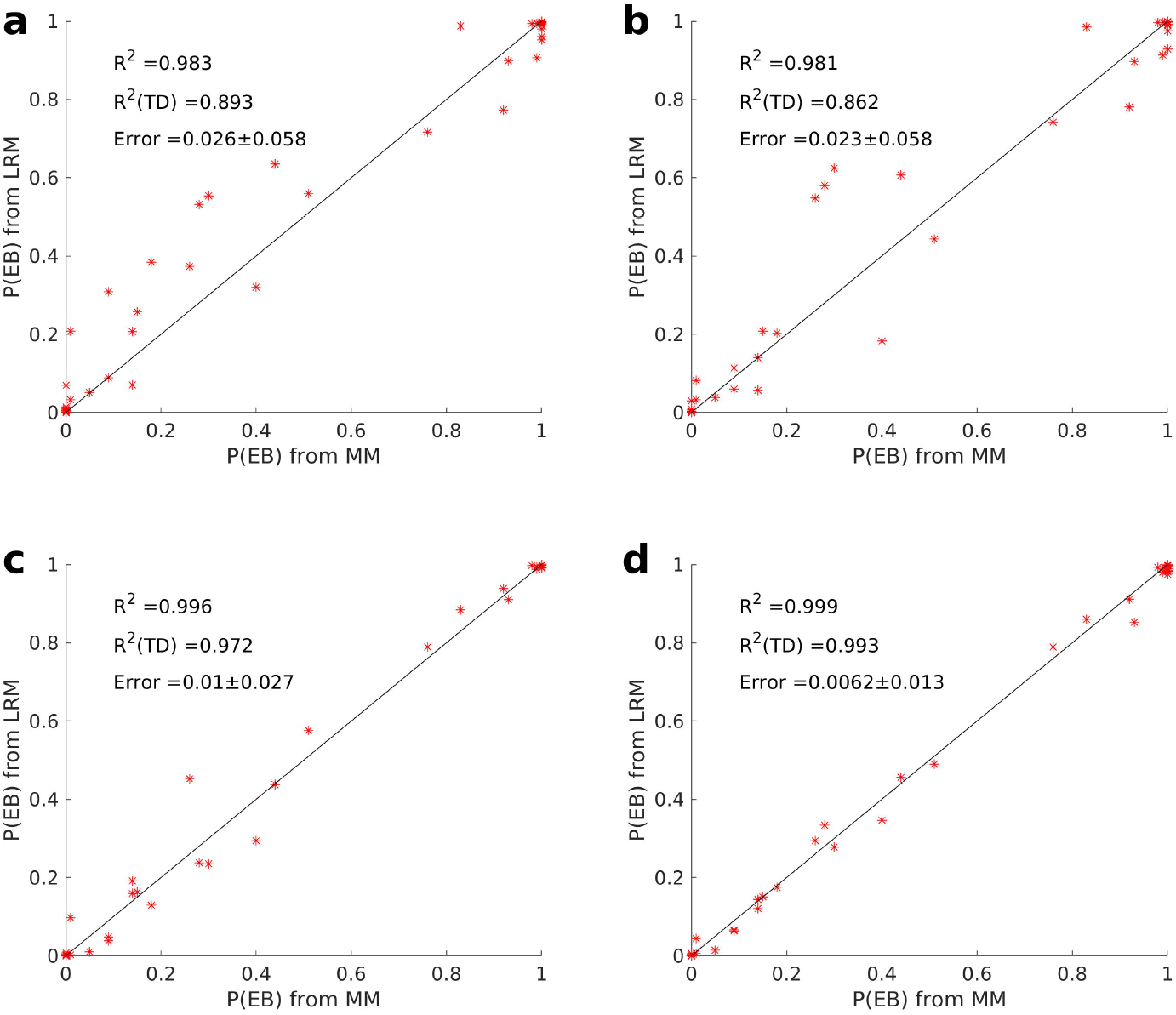
DAD-induced ectopic beat (EB) prediction performance for different LRMs. Performance of each LRM is evaluated on the same test set which consists of 100 myocyte model input (MMI) sets for all models. Each LRM has a unique training strategy. A) LRM trained with only linear features (excluding quadratic features) and only 100 MMI sets from only the first iteration. B) LRM trained with only linear features (excluding quadratic features) and 200 MMI sets (100 sets in the first iteration + 100 transition domain sets in the second iteration). C) LRM trained with linear features and quadratic features on 100 MMI sets from only the first iteration. D) Reproduction of Fig. 1E, LRM trained with linear features and quadratic features and 200 MMI sets (100 sets in the first iteration + 100 transition domain sets in the second iteration). Linear features and quadratic features used in A-D are given in Table S4. First iteration training data was identical for all LRMs and second iteration training data was identical for LRMs in panels B and D. R^2^ for the entire region of interest and transition domain as well as the average error are reported.

**Fig. S3.**
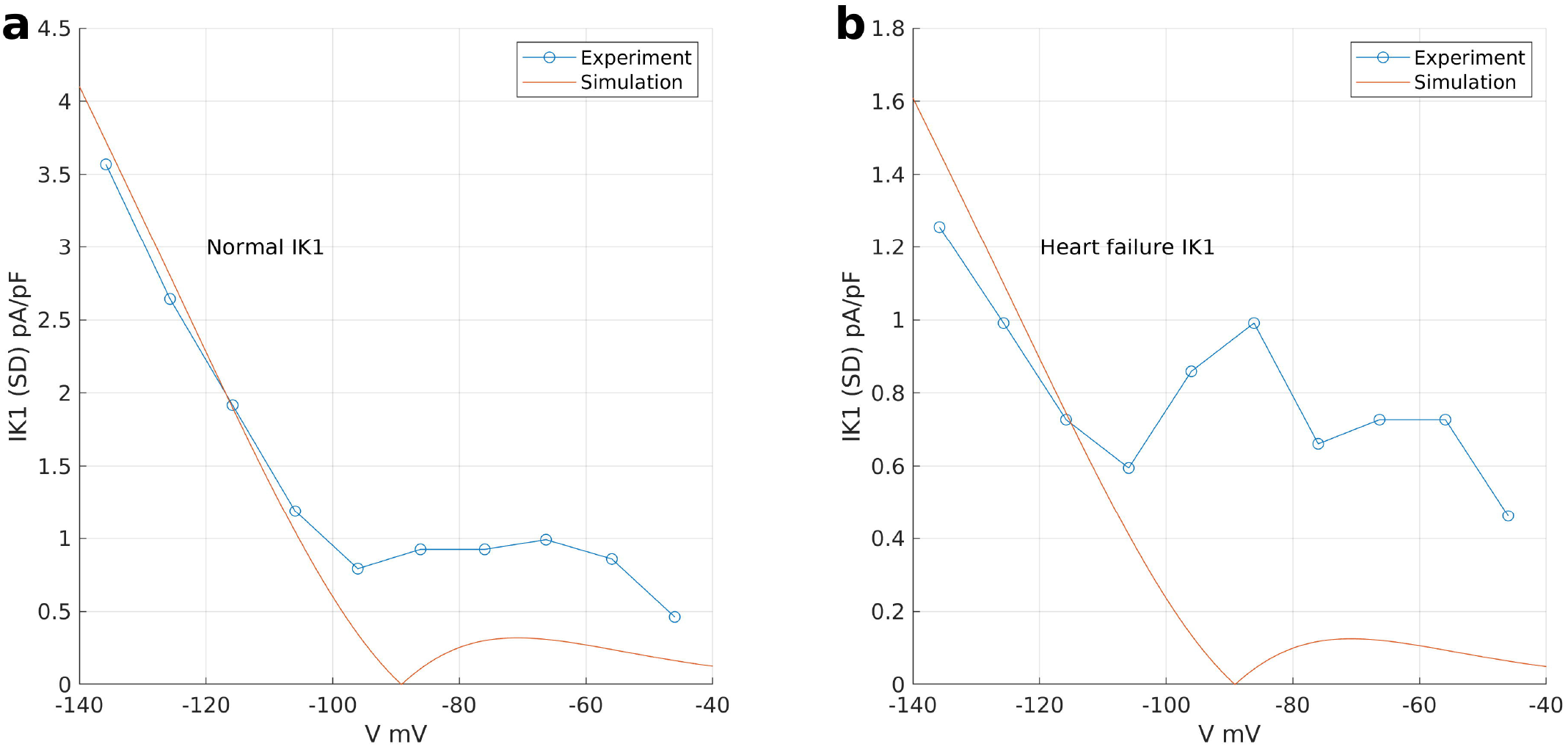
Constraining G_K1_ scaling factor (G_K1__sf). Experimental I_K1_ I-V relationship for normal and Heart failure (HF) is measured and showed in Fig. 5C of Pogwizd et. al. In the experimental protocol, I_K1_ is measured in response to 500-ms steps from a holding potential of -30 mV to test potentials in the range of -120 mV to +40 mV in 6 normal cells and 6 HF cells. The detail of the protocol is described in Pogwizd et. al. Error bars represent standard error (SE). (A) Best fit of the standard deviation (SD) for the normal I_K1_ using the same voltage clamp protocol. (B) Best fit of the SD for the HF I_K1_ using the same voltage clamp protocol.

**S1 Table.** MMIs chosen for P(EB) prediction.

| MMIs | Lower bound | Upper bound |
| --- | --- | --- |
| **Initial $[Ca]_i$ (nM)** | 100 | 300 |
| **Initial $[Ca]_{SR}$ (mM)** | 0.3 | 0.7 |
| **$G_{K1\_sf}$** | 0 | 1 |
| **$k_{RyR^+\_sf}$** | 2.5 | 7.5 |

**S1 Text. Supporting description of logistic regression model and entropy.**

