## Supplementary material for "Estimating Ectopic Beat Probability with Simplified Statistical Models that Account for Experimental Uncertainty": S1 Text: Supporting description of logistic regression model and entropy

### Supporting Information

#### S1 Text. Supporting description of logistic regression model and emtropy.

**Logistic regression model**. S1 Fig shows the workflow for building a two-iteration logistic regression model. In the first iteration, the first step is to identify M MMIs: $MMI_i$, i = 1, 2, …, M-1, M. Four MMIs are chosen for the EB study as described in the Method. The region of interest (ROI) for each MMI, representing the biophysical meaningful range of the values of $MMI_i$, is then determined based on experiments and defined by an upper bound ($UB_i$) and a lower bound ($LB_i$). The ROI of the full set of MMIs makes up a M-dimensional MMI space $\Omega \in R^M$. We randomly generate N MMI sets (MMISets) $MMISet_j \in \Omega$, j = 1, 2, …, N with a uniform distribution. The $j^{th}$ MMI set $MMISet_j$ contains M MMI values, where $MMISet_j = [mmi_{1j}, mmi_{2j}, …, mmi_{Mj}]^T$. Stochastic myocyte model (MM) simulations are performed L times for each $MMISet_j$ to obtain L realizations per MMI set. The detailed MM simulation protocol is described in Methods. The output of the MM for each $MMISet_j$ will be a vector of length L with binary elements where each element indicates whether (= 1) or not (= 0) the event of interest (in this case EAD or EB) occurred. A probability of the event occurrence, $P_j$(event), is estimated from the output vector for each $MMISet_j$. Then, we performed logistic regression on all MMI sets and their corresponding event probabilities. More specifically, the MxN feature matrix is [$MMISet_1$, $MMISet_2$, …, $MMISet_N$] and the 1xN label vector is [$P_1$(event), $P_2$(event), …, $P_N$(event)]. The logistic equation takes the form

$$P(event)=\frac{1}{1+\exp(-\underline{B}_o^T \underline{P}_o)} \quad (S1)$$

, where $\underline{P}_o$ = [$MMI_1$, $MMI_2$,…, $MMI_M$] is the feature vector input for the logistic equation, $\underline{B}_o = [b_0, b_1,…, b_M]$ is the vector of weights and P(event) is the predicted event probability from the logistic equation. In summary, in this first iteration (step 1 – 6 in S1 Fig), we perform a logistic regression on the feature matrix and the label vector, and this yields our first estimation of logistic equation weights ($\underline{B}_o$).

After the first iteration, we use $\underline{B}_o$ with S1 Eq to create an additional constraint in addition to that imposed by the ROI in step 2 of S1 Fig to estimate the transition domain (TD)

$$P(event)_{LB} < P(event)=\frac{1}{1+\exp(-\underline{B}_o^T \underline{P}_o)} < P(event)_{UB} \quad (S2)$$

where $P(event)_{LB}$ is the lower bound for P(event) and $P(event)_{UB}$ is the upper bound for P(event). In this study, we chose $P(event)_{LB}$ = 0.01 and $P(event)_{UB}$ = 0.99. We repeat steps 3, 4, and 5 described in S1 Fig on the estimated TD from S2 Eq. Therefore, we obtain an additional N newly generated MMISets. Combining with the MMISets generated in the first iteration, the new feature matrix will then be [$MMISet_1$, …, $MMISet_N$, $MMISet_{N+1}$, …, $MMISet_{2N}$]. Label vectors becomes [$P_1$(event), …,

$P_N$(event), $P_{N+1}$(event), …, $P_{2N}$(event)]. In order to further improve the model performance, we derived quadratic features from the MMIs in the form of $MMI_{i1}$*$MMI_{i2}$, i1, i2 = 1, 2, …, N. Besides linear features (MMIs), we enumerated all possible combinations of quadratic features (Qs) and choose those with the highest consistent Akaike information criterion (CAIC) [1] for our model. After adding H selected quadratic terms into feature matrix, we conduct logistic regression again with the updated feature matrix and label vector to obtain the second iteration logistic equation

$$\text{P(event)}=\frac{1}{1+\exp(-\underline{B}_f^T \underline{P}_f)} \tag{S3}$$

where $\underline{P}_f$ = [$MMI_1$, $MMI_2$, …, $MMI_M$, $Q_1$, $Q_2$, …, $Q_H$] is the feature vector ($Q_1$, …, $Q_H$ are quadratic features) and $\underline{B}_f$ = [$b_0$, $b_1$,…, $b_{M+H}$] is the vector of weights. S3 Eq is our final simplified model which is referred to simply as the logistic regression model (LRM).

**Entropy.** Entropy, a measure of the uncertainty of random variables (RVs) [2], is more appropriate than the variance to assess uncertainty for multimodal distributions [3]. The RV with larger uncertainty will have a larger entropy. Given the fact that the P(EB) distribution can become bimodal, we use the entropy to quantify P(EB) uncertainty. We separated the P(EB) distributions into 100 bins where the interval for *i*th bin is [0.01*i, 0.01*i+0.01], i=0, 1, …, 99. Assume that the *i*th bin's sample frequency is $freq_i$, the entropy of P(EB) distribution is

$$Entropy = -\sum_{i=0}^{99} freq_i \ln(freq_i) \tag{S4}$$
